# NeuroGFX: a graphical functional explorer for fruit fly brain circuits

**DOI:** 10.1101/092437

**Authors:** Chung-Heng Yeh, Yiyin Zhou, Nikul H. Ukani, Aurel A. Lazar

## Abstract

Recently, multiple focused efforts have resulted in substantial increase in the availability of connectome data in the fruit fly brain. Elucidating neural circuit function from such structural data calls for a scalable computational modeling methodology. We propose such a methodology that includes i) a brain emulation engine, with an architecture that can tackle the complexity of whole brain modeling, ii) a database that supports tight integration of biological and modeling data along with support for domain specific queries and circuit transformations, and iii) a graphical interface that allows for total flexibility in configuring neural circuits and visualizing run-time results, both anchored on model abstractions closely reflecting biological structure. Towards the realization of such a methodology, we have developed NeuroGFX and integrated it into the architecture of the Fruit Fly Brain Observatory (http://fruitflybrain.org). The computational infrastructure in NeuroGFX is provided by Neurokernel, an open source platform for the emulation of the fruit fly brain, and NeuroArch, a database for querying and executing fruit fly brain circuits. The integration of the two enables the algorithmic construction/manipulation/revision of executable circuits on multiple levels of abstraction of the same model organism. The power of this computational infrastructure can be leveraged through an intuitive graphical interface that allows visualizing execution results in the context of biological structure. This provides an environment where computational researchers can present configurable, executable neural circuits, and experimental scientists can easily explore circuit structure and function ultimately leading to biological validation. With these capabilities, NeuroGFX enables the exploration of function from circuit structure at whole brain, neuropil, and local circuit level of abstraction. By allowing for independently developed models to be integrated at the architectural level, NeuroGFX provides an open plug and play, collaborative environment for whole brain computational modeling of the fruit fly.

## Additional Details

NeuroGFX builds upon two major modules of the FFBO architecture [1], Neurokernel [2] and NeuroArch [3]. Both have been designed with the explicit aim of facilitating the generation of executable neural circuit models on GPUs. NeuroGFX conjoins the simultaneous operations of model exploration on NeuroArch and model execution on Neurokernel in a unified graphical web interface, presents queried connectomic data as a reconfigurable circuit diagram, and renders interactive simulation results. NeuroGFX facilitates construction and execution of neural circuitry at various brain levels of abstraction. We demonstrate below the capabilities of NeuroGFX on the whole brain, neuropil and local circuit level.

## NeuroGFX whole brain level exploration

The fruit fly brain is decomposed into some 50 neuropils. Based on [4], we laid out the circuit diagram of the network of Local Processing Units, i.e., model abstractions of neuropils. In Figure 1, the circuit diagram on the brain level is shown side-by-side with the anatomical representation of neuropils (blue) and tracts (bright-colors). Through the interface, the circuit diagram can be reconfigured to include any subset of neuropils. On the bottom of Figure 1 only the neuropils of the early vision system and their interconnects are shown. Circuit models of neuropils and their interconnects can be subsequently retrieved from NeuroArch and be explored through execution in Neurokernel. Although it is still far from having the needed data to execute the entire fly brain, NeuroGFX on the whole brain level lays out the guidelines for the development of whole brain emulation, allowing us to elucidate the functional role of individual neuropils in a subsystem.

**Figure 1:**
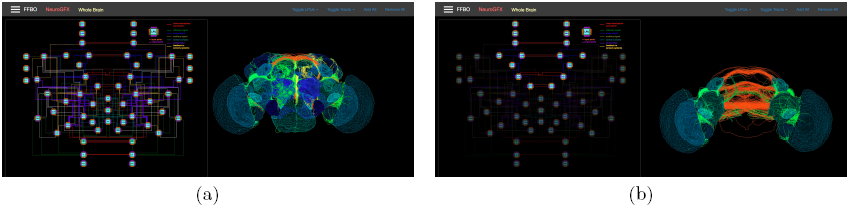
(a) Whole brain circuit diagram and anatomy. (b) Circuit diagram and anatomy of the vision system.

## NeuroGFX neuropil level exploration

The antennal lobe (AL) is the first neuropil of the olfactory system in the fly brain, consisting of some 50 subregions called glomeruli. In Figure 2, we reconstruct the antennal lobe to interrogate its functionality, and visualize the structure of glomeruli along with the reconfigurable circuit diagram of the antennal lobe. The effect of pathological synapse activity in the antennal lobe is evaluated by increasing the synaptic strength of the inhibitory local neurons. In the same figure, the simulation results of healthy and diseased circuits are presented in both biological view (upper) and interactive plots (lower).

**Figure 2:**
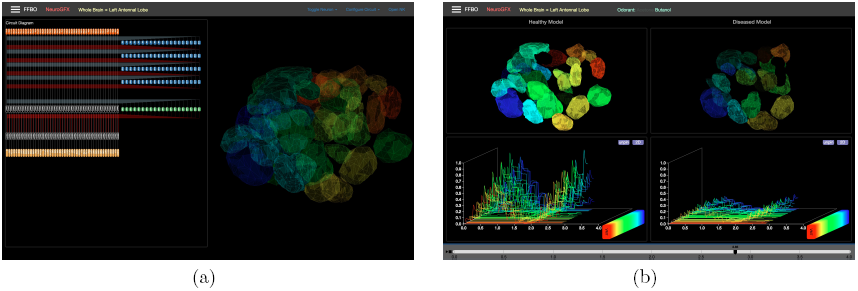
(a) Antennal lobe circuit diagram and anatomical visualization. (b) Comparison of simulation results between healthy and diseased models.

## NeuroGFX local circuits level exploration

The lamina is the first neuropil of the early visual system of the fruit fly, receiving inputs directly from the retina. The retinotopic map is preserved in the lamina by columnar circuits, called cartridges. We use NeuroGFX to explore the function of these local circuits. The complete shape of neurons and connectivity between neurons in a single cartridge of the lamina have been determined by serial electron microscopy [5] Using NeuroGFX, we visualize both the morphology (skeletons) of the neuron and the circuit diagram side by side (see Figure 3 top). By clicking on the Load Cartridge button, NeuroGFX is instructed to fire a series of queries to NeuroArch database where a model of the retina and lamina network resides. Upon retrieving the circuit model, information about individual neuron can be shown by hovering the mouse over the neuron in the circuit diagram. Similar to the case of the brain and neuropil level, NeuroGFX enables users to add/remove any neurons in the circuit diagram. The configured circuit is then sent to Neurokernel for execution. The inputs to each of the photoreceptors are rendered on the top of the result viewer while the response of all neurons are shown on the bottom (see Figure 3 bottom).

**Figure 3:**
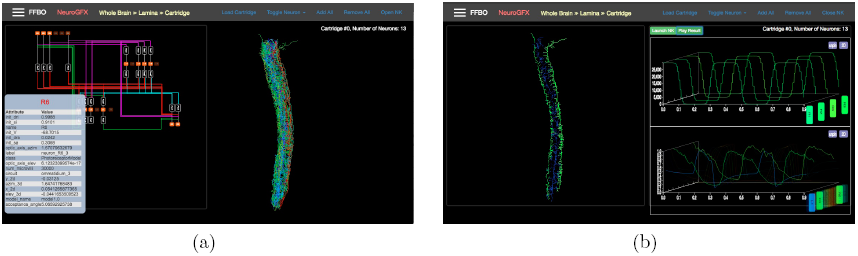
(a) Data retrieval of a lamina cartridge model and executable circuit construction using NeuroGFX. (b) Results of circuit execution.

## References

[1] N.H. Ukani et al., The Fruit Fly Brain Observatory: from structure to function, bioRxiv, 092288, 2016. DOI: 10.1011/092288.

[2] L.E. Givon and A.A. Lazar, PLoS ONE 11(1): e0146581, 2016. DOI: 10.1371/journal.pone.0146581.

[3] L.E. Givon et al., NeuroArch: A Graph dB for Querying and Executing Fruit Fly Brain Circuits, Neurokernel Request for Comments, Neurokernel RFC#4, 2015. DOI: 10.5281/zen-odo.44225

[4] C.-T. Shih et al., Curr. Biol 25(10): 1249–1258 2015, DOI: 10.1016/j.cub.2015.03.021

[5] M. Rivera-Alba et al., Curr. Biol 21(23): 2000–2005, 2011, DOI: 10.1016/j.cub.2011.10.022

